# Altered Dynamic Functional Connectivity of the Frontoparietal Network in Major Depressive Disorder: Evidence from a Large-Scale Resting-State fMRI Study

**DOI:** 10.1101/2025.09.17.676759

**Authors:** Parsa Seyedzadeh, Seyed Armin Hosseini

## Abstract

Major depressive disorder (MDD) is a prevalent psychiatric condition characterized by affective disturbances and cognitive deficits. Among these, cognitive inflexibility and executive dysfunction are particularly prominent, yet the temporal dynamics of the frontoparietal control network (FPN), a core substrate of cognitive control, remain poorly understood. Using harmonized resting-state fMRI data from the REST-meta-MDD consortium (n = 887; 442 MDD, 445 healthy controls), we investigated dynamic functional connectivity (dFC) within the FPN. Time-varying correlations among 21 FPN nodes were estimated using a sliding-window approach and clustered via k-means to identify recurring connectivity states. Temporal metrics included fractional occupancy, mean dwell time, and transition counts. Three unique FPN states were recognized. In comparison to healthy individuals, those with Major Depressive Disorder (MDD) exhibited prolonged durations in a hypoconnected state, extended dwell times in this configuration, and fewer total transitions, indicating diminished neural flexibility. Direct transitions between low-connectivity (hypoconnected) and high-connectivity (hyperconnected) states were selectively diminished, indicating a disruption in the direct transition between two functionally distinct states of the FPN. Overall, these findings reveal a fundamental disruption in the temporal organization of frontoparietal connectivity in MDD, marked by predominant hypoconnectivity, reduced flexibility, and constrained state transitions. By delineating the dynamic properties of network function, this study advances a mechanistic framework for interpreting prior inconsistencies in static connectivity research and underscores the necessity of time-resolved approaches in characterizing large-scale network dysfunction in psychiatric disorders.

## 1 INTRODUCTION

Major Depressive Disorder (MDD) is a prevalent and debilitating psychiatric disorder impacting over 270 million individuals globally, acknowledged as a primary contributor to non-fatal health loss, with more than 37 million years lived with disability (YLDs) attributed to it in 2019 (Yan et al., 2024). Clinically, Major Depressive Disorder (MDD) includes a constellation of emotional symptoms, notably enduring sorrow and anhedonia, alongside cognitive deficits in executive function, attention, inhibitory control, and emotion regulation. These cognitive abnormalities contribute to an elevated risk of recurrence, sustained impairment, and unfavorable functional outcomes (Buckner et al., 2008; Jarrett et al, 2012; Joormann & Vanderlind, 2014; Paykel, 2008; Tian et al., 2016). Nevertheless, diagnosis continues to rely on symptom-based categories such as the DSM, which do not incorporate biological characteristics, underscoring the necessity for neurobiological markers that can offer objective and quantifiable indicators of disease state, severity, and progression (Young et al., 2016).

Due to the necessity for objective biomarkers, resting-state functional magnetic resonance imaging (rs-fMRI) has arisen as a method for examining the neurological framework of major depressive disorder (MDD). Functional connectivity analyses of resting-state fMRI data indicate that Major Depressive Disorder (MDD) is linked to impaired communication among extensive brain networks, notably the default mode network (DMN), salience network (SN), and frontoparietal control network (FPN), which likely contribute to cognitive and emotional deficits associated with this condition (Fu-Cai, 2024; Dichter et al., 2015; Ma et al., 2024; Zhang et al., 2025; Demirtas et al., 2016; Wei et al., 2013; Kaiser et al., 2015; Balzekas et al., 2018; Zhang et al., 2018).

The frontoparietal control network (FPN) is an extensive system situated in the lateral prefrontal and posterior parietal cortices, serving as a versatile hub for the coordination of distributed brain activity (Cole et al., 2013; Power et al., 2011). This network incorporates cognitive and emotional processes vital for adaptive behavior through its important function in top-down cognitive control, attention distribution, and emotion regulation (Buhle et al., 2014; Dosenbach et al., 2008; Gao et al., 2024; Wu et al., 2024). In Major Depressive Disorder (MDD), alterations in the fronto-parietal network (FPN) connectivity have been consistently linked to deficits in executive function, inhibitory control, and attention management (Yu et al., 2019; Schultz et al., 2017).

Increased empirical data underscores the importance of FPN deficiency in depression. Irregular connections within this network have been associated with maladaptive self-referential thinking and rumination, resulting in the exacerbation of negative emotion and cognitive rigidity (Doston et al., 2020; Rai et al., 2021; Li et al., 2020; Wagner et al., 2016; Yang et al., 2024; Zheng et al., 2024). Collectively, these converging findings indicate that the FPN serves as a pivotal neuronal substrate for the cognitive-emotional phenomenology of MDD and constitutes a feasible target for biomarker development.

Despite accumulating evidence concerning FPN disruption in MDD, the exact nature of within-network alterations remains ambiguous and inconsistent. Some studies have indicated diminished intrinsic connection (hypoconnectivity) in the FPN, signifying a reduction in cognitive control capacity. Meta-analytic studies and empirical research have demonstrated a considerable decrease in within-FPN connectivity in rs-fMRI data (Hwang et al., 2015; Lan et al., 2022; Machaj et al., 2024).

Conversely, certain extensive investigations have not detected significant disparities in FPN connectivity between depressed and healthy persons, nor have they identified substantial within-network alterations in the FPN (Javaheripour et al., 2021; Whitton et al., 2018). Recently, contradictory evidence has surfaced. Extensive meta-analyses utilizing rs-fMRI and electrophysiological studies have indicated heightened intrinsic connectivity within the FPN in depression, with certain research associating excessive connectivity in this network with severe depressive symptoms, including suicide risk (Zhang et al., 2025; Ren et al., 2025; Chu et al., 2025). These contradictions hinder the elucidation of how FPN dysfunction might explain the cognitive and executive deficiencies in depression, hence diminishing its reliability as a brain marker.

One of the possible explanations for this inconsistency can be the restriction of studies to static functional connectivity and neglecting the temporal variability in coupling within this network, which treats connectivity as stable across the entire scan. This assumption neglects the inherently dynamic nature of brain network interactions. A growing body of research shows that functional connectivity fluctuates over time, even during rest, and that these temporal dynamics are relevant to cognition and psychiatric symptoms (Demirtas et al., 2016; Marchitelli et al., 2022).

Beyond the limitations of static connectivity approaches, inconsistencies across prior studies may also reflect substantial heterogeneity in clinical characteristics, genetic and environmental factors, and analytic strategies. Small, single-site datasets exacerbate these issues by limiting statistical power and generalizability, increasing the risk of spurious findings (Chen et al., 2022; Yu et al., 2018). Large, harmonized multisite datasets are therefore essential for reducing site-related biases and achieving more robust estimates of group-level effects.

To address these gaps, the present study investigates the dynamic properties of frontoparietal network (FPN) connectivity in major depressive disorder (MDD) using resting-state fMRI data from the REST-meta-MDD consortium, the largest coordinated dataset available for this disorder (Yan et al., 2019). Dynamic functional connectivity (dFC) metrics, fractional occupancy, mean dwell time, and number of state transitions were derived to capture temporal stability and flexibility of the FPN, processes theoretically linked to cognitive inflexibility and executive dysfunction in MDD.

Using a sliding-window correlation approach applied to 21 FPN nodes, followed by k-means clustering of windowed connectivity patterns, we identified recurring network states and extracted temporal features for comparison between patients with MDD and healthy controls. This study thus provides the first large-scale analysis of FPN dFC in depression, aiming to resolve prior inconsistencies and evaluate whether altered temporal properties of frontoparietal connectivity represent a reliable network-level signature of the disorder.

## 2 METHODS

### 2.1 Participants and Dataset

Data for this study were obtained from the REST-meta-MDD project, the largest coordinated effort to collect rs-fMRI data in MDD to date (Yan et al., 2019). The full consortium dataset includes 2,428 participants (1,300 patients with MDD and 1,128 healthy controls, HCs) recruited by 17 research groups across 25 hospitals in China.

For the present analysis, we focused on participants who met standardized inclusion criteria to ensure both clinical validity and data quality. Eligible individuals were between 18 and 65 years of age and had at least 5 years of formal education. Patients were diagnosed with MDD according to DSM-IV criteria (First et al., 1997), confirmed through structured clinical interviews conducted by trained psychiatrists, and were required to have a score of at least 8 on the 17-item Hamilton Depression Rating Scale (HAMD-17; Williams, 1988) at the time of scanning. To ensure robust estimation of dFC, only sites with resting-state scans lasting at least 8 minutes (TR = 2 seconds, ≥240 volumes) were included, consistent with prior work showing that longer scan lengths yield more reliable estimates of time-resolved connectivity (Birn et al., 2013). Functional data were parcellated using the Dosenbach 160-region atlas (Dosenbach et al., 2010), a well-validated set of regions of interest optimized for connectivity analyses.

Participants were excluded if their behavioral or imaging data were incomplete or if their functional scans showed missing signal across multiple brain regions. After applying these criteria, the final dataset comprised 442 patients with MDD and 445 HCs collected across seven sites. Demographic and clinical characteristics of the included participants are summarized in Table 2, with site-specific distributions presented in Table 1. Group differences in demographic measures, including age, sex distribution, HAMD scores, and illness duration, were examined using Welch’s two-sample *t*-tests for age and *χ*^2^ tests for sex. No significant differences were observed between the age and sex. All study protocols were approved by the institutional review boards of the participating sites, and written informed consent was obtained from all participants in accordance with the Declaration of Helsinki.

**TABLE 1:** Site-specific demographic and clinical characteristics of healthy controls (HCs) and patients with major depressive disorder (MDD). Data are presented as mean *±* standard deviation where applicable. HAMD = Hamilton Depression Rating Scale.

| Site | Variable | HC | MDD |
| --- | --- | --- | --- |
| 15 | Number of subjects | 36 | 35 |
| | Age (years) | 39.19 ± 14.46 | 46.80 ± 12.84 |
|  | Sex (M/F) | 17 / 19 | 11 / 24 |
| | HAMD | – | 26.43 ± 5.11 |
| | Illness duration (months) | – | 34.47 ± 53.81 |
| 17 | Number of subjects | 42 | 31 |
| | Age (years) | 21.57 ± 6.32 | 21.84 ± 3.16 |
|  | Sex (M/F) | 13 / 29 | 7 / 24 |
| | HAMD | – | 19.52 ± 5.30 |
|  | Illness duration (months) | – | – |
| 19 | Number of subjects | 30 | 24 |
| | Age (years) | 35.63 ± 10.09 | 39.33 ± 11.43 |
|  | Sex (M/F) | 14 / 16 | 4 / 20 |
| | HAMD | – | 21.62 ± 6.62 |
| | Illness duration (months) | – | 90.75 ± 99.35 |
| 20 | Number of subjects | 229 | 235 |
| | Age (years) | 39.57 ± 15.74 | 38.46 ± 11.93 |
|  | Sex (M/F) | 73 / 156 | 81 / 154 |
| | HAMD | – | 21.43 ± 5.05 |
| | Illness duration (months) | – | 50.09 ± 65.53 |
| 21 | Number of subjects | 65 | 64 |
| | Age (years) | 36.51 ± 12.52 | 34.47 ± 13.15 |
|  | Sex (M/F) | 28 / 37 | 32 / 32 |
| | HAMD | – | 17.58 ± 6.41 |
| | Illness duration (months) | – | 90.46 ± 99.47 |
| 22 | Number of subjects | 20 | 29 |
| | Age (years) | 24.35 ± 7.07 | 34.14 ± 9.95 |
|  | Sex (M/F) | 12 / 8 | 14 / 15 |
| | HAMD | – | 22.79 ± 5.02 |
| | Illness duration (months) | – | 38.90 ± 43.63 |
| 23 | Number of subjects | 23 | 24 |
| | Age (years) | 33.00 ± 12.05 | 26.00 ± 7.96 |
|  | Sex (M/F) | 8 / 15 | 14 / 10 |
| | HAMD | – | 21.58 ± 5.36 |
| | Illness duration (months) | – | 25.21 ± 29.12 |

**TABLE 2:** Overall demographic and clinical characteristics of patients with major depressive disorder (MDD) and healthy controls (HCs). Data are presented as mean *±* standard deviation where applicable. Group differences were assessed using Welch’s *t*-tests (for age) or *χ*^2^ tests (for sex). HAMD = Hamilton Depression Rating Scale.

| Variable | HC | MDD | df | t | $\chi^2$ | p |
| --- | --- | --- | --- | --- | --- | --- |
| Number of subjects | 445 | 442 | – | – | – | – |
| Age (years) | 36.11 ± 14.82 | 36.47 ± 12.77 | 867.7 | 0.38 | – | 0.698 |
| Sex (M/F) | 165 / 280 | 163 / 279 | 1.0 | – | 0.00 | 1.000 |
| HAMD | – | 21.24 ± 5.76 | – | – | – | – |
| Illness duration (months) | – | 55.65 ± 73.49 | – | – | – | – |

### 2.2 Data Preprocessing

rs-fMRI data were preprocessed using the Data Processing Assistant for rs-fMRI (DPARSF; Yan & Zang, 2010) following the standardized protocol of Yan et al. (2019). The first 10 volumes were removed, after which slice-timing correction, realignment, and co-registration of each participant’s T1-weighted image to the mean functional image were performed. Structural images were segmented into gray matter, white matter, and cerebrospinal fluid, and functional data were normalized to Montreal Neurological Institute (MNI) space and resampled to 3 mm isotropic voxels.

Nuisance signals, including the Friston 24 head-motion parameters, white matter, cerebrospinal fluid, and the global signal, were regressed out, and a linear trend term was included to control for scanner drifts. Participants with a mean framewise displacement greater than 0.2 mm were excluded. Finally, temporal band-pass filtering (0.01–0.1 Hz) was applied to the time series to retain frequencies relevant to resting-state activity.

### 2.3 Dynamic Functional Connectivity Analysis

dFC was estimated using the Dynamic Brain Connectome toolbox (DynamicBC, version 2.2; Liao et al., 2014) implemented in MATLAB 2022 (The MathWorks, 2022). Regional BOLD time series were extracted from the 21 nodes of the FPN defined by the Dosenbach 160 atlas (Dosenbach et al., 2010). To capture time-varying changes in connectivity, we employed a sliding-window correlation approach. Each participant’s time series was segmented into windows of 22 TRs (44 s) with a step size of 1 TR (2 s), yielding 209 overlapping windows across the full resting-state scan. These parameters were chosen based on the work of Allen et al. (2014), which demonstrated their effectiveness in capturing reliable fluctuations in resting-state dynamic connectivity while providing a reasonable balance between temporal resolution and the stability of correlation estimates. Within each window, Pearson correlation coefficients were computed between all pairs of FPN nodes, resulting in a 21 × 21 connectivity matrix. The correlation values were then Fisher z-transformed to improve the normality of the distribution (Figure 1).

**FIGURE 1:**
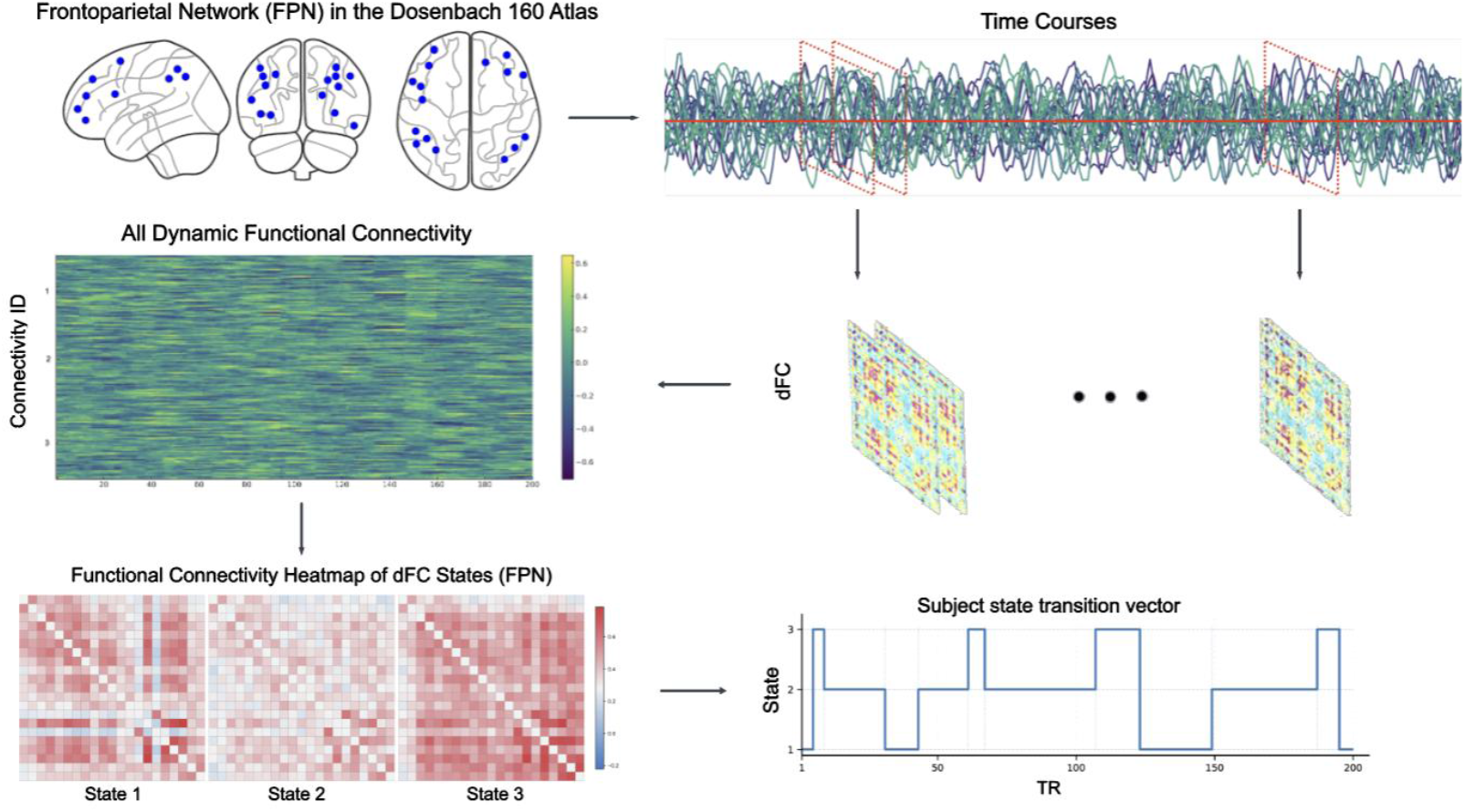
Analysis pipeline for dynamic frontoparietal network (FPN) connectivity. BOLD time series were extracted from 21 FPN nodes defined by the Dosenbach 160 atlas (top left). A sliding window (22 TR; step = 1 TR) was applied to each subject’s time courses (top right), and Fisher *z*–transformed Pearson correlations were computed within each window to yield a sequence of 21 *×* 21 dynamic functional connectivity (dFC) matrices (middle). All windowed matrices across participants were concatenated and clustered with *k*-means (*k* = 3) to identify recurring FPN states (bottom left; state centroids shown as connectivity heatmaps). For each participant, the resulting state sequence (bottom right) was used to compute temporal features: fractional occupancy, mean dwell time, and total number of state transitions.

Following the construction of windowed matrices, additional preprocessing was performed in Python 3.10 (Python Software Foundation, 2022) to prepare the data for subsequent analyses. Correlation values were numerically stabilized by shifting boundary values (±1) by a small epsilon to prevent infinities, after which Fisher’s r-to-z transformation was applied. The upper-triangular elements of each matrix (excluding the diagonal) were then extracted to form feature vectors representing the connectivity patterns within each time window. Matrices containing invalid values (e.g., NaNs or infinities) were excluded to ensure data quality.

This procedure yielded a set of time-resolved connectivity vectors for each participant, characterizing the evolving temporal organization of FPN connectivity across the scan. These dynamic connectivity vectors were concatenated across all participants and used as input for the subsequent group-level k-means clustering analysis.

An overview of the FPN dFC pipeline is shown in Figure 1: we extracted BOLD time series from 21 FPN nodes, applied a sliding-window correlation (22 TR, step 1 TR) to obtain windowed 21×21 matrices, and then clustered the vectorized windows; the resulting state sequence per subject yielded fractional occupancy, mean dwell time, and transition counts.

### 2.4 Clustering Analysis

To identify recurring patterns of dFC within the FPN, we applied k-means clustering to the windowed connectivity vectors derived from the 21 FPN nodes of the Dosenbach 160 atlas (Dosenbach et al., 2010). Clustering was performed with the Euclidean (L2) distance metric, a standard approach in dFC research, which produced the same results as the Manhattan (L1) distance in our high-dimensional setting while being more computationally efficient (Allen et al., 2014).

To enhance stability, we first identified exemplar windows, defined as local maxima in the temporal variance of connectivity across ROI pairs. Exemplar windows from all participants were pooled and clustered with k-means using 500 random initializations (replicates) and allowing up to 500 iterations per run. These group-level centroids were then used to initialize a second round of k-means clustering applied to the full set of FPN dFC windows across all subjects, minimizing sensitivity to random starting conditions. The optimal number of clusters was determined using the elbow criterion (Tibshirani et al., 2001), which showed an inflection at *k* = 3 (Figure 2).

**FIGURE 2:**
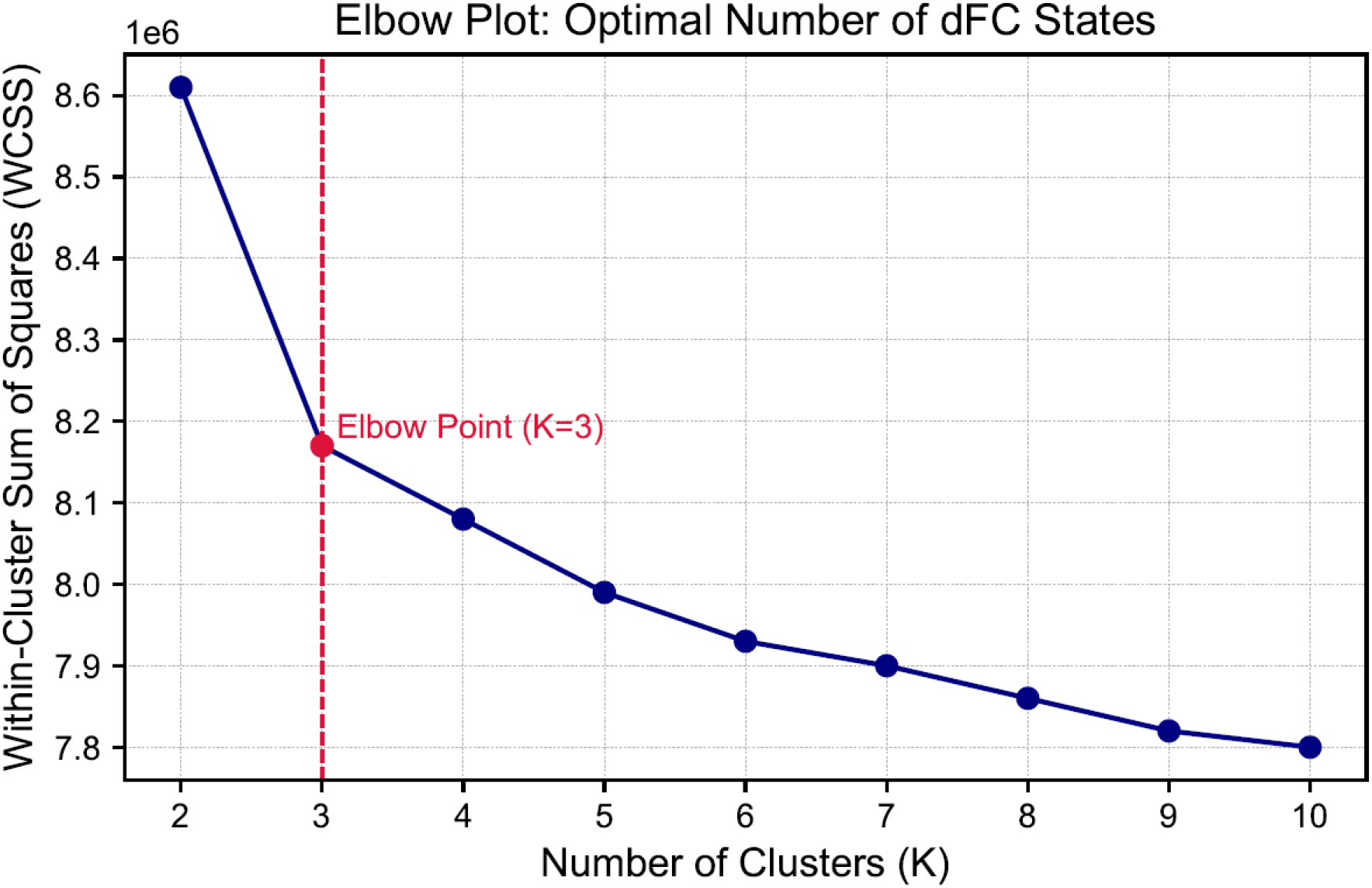
Elbow plot for determining the optimal number of clusters (*k*) in the *k*-means clustering of dynamic functional connectivity (dFC) states within the frontoparietal network (FPN). The plot shows within-cluster sum of squares (WCSS) as a function of *k*. The inflection point at *k* = 3 indicates the optimal balance between model complexity and within-cluster homogeneity.

As shown in Figure 2, the elbow at three clusters indicated a balance between parsimony and within-cluster homogeneity. Accordingly, each dFC window was assigned to a connectivity state based on the minimum Euclidean distance between its vectorized connectivity pattern and the final cluster centroids. From these state sequences, we derived three temporal features to characterize individual dFC dynamics. Fractional occupancy was defined as the proportion of windows assigned to a given state, indexing the predominance of that connectivity configuration. Mean dwell time measures the average number of consecutive windows spent in a state before switching, reflecting temporal stability.

The number of transitions captured the total frequency of state changes across the scan, providing an index of network flexibility.

### 2.5 Statistical Analysis

#### 2.5.1 Within-subject comparison of state-wise FC strength

To verify that the three connectivity states identified through clustering represented meaningfully distinct patterns of frontoparietal functional coupling, we compared mean FC strength across states within individuals. For each participant, FC strength was defined as the average of Fisher *z*-transformed pairwise correlations among the 21 FPN nodes. A repeated-measures ANOVA was applied with state as the within-subject factor. When the assumption of sphericity was violated, the Greenhouse–Geisser correction was used to adjust the degrees of freedom. Significant omnibus effects were followed by paired post-hoc *t*-tests, with *p*-values corrected for multiple comparisons using the Benjamini–Hochberg false discovery rate (FDR; *α* = 0.05). Notably, nearly all participants from both groups occupied each state at least once during the resting-state scan Figure 3, ensuring that the subsequent within-subject comparisons were based on nearly complete samples and that the identified states reflect robust and generalizable connectivity configurations.

**FIGURE 3:**
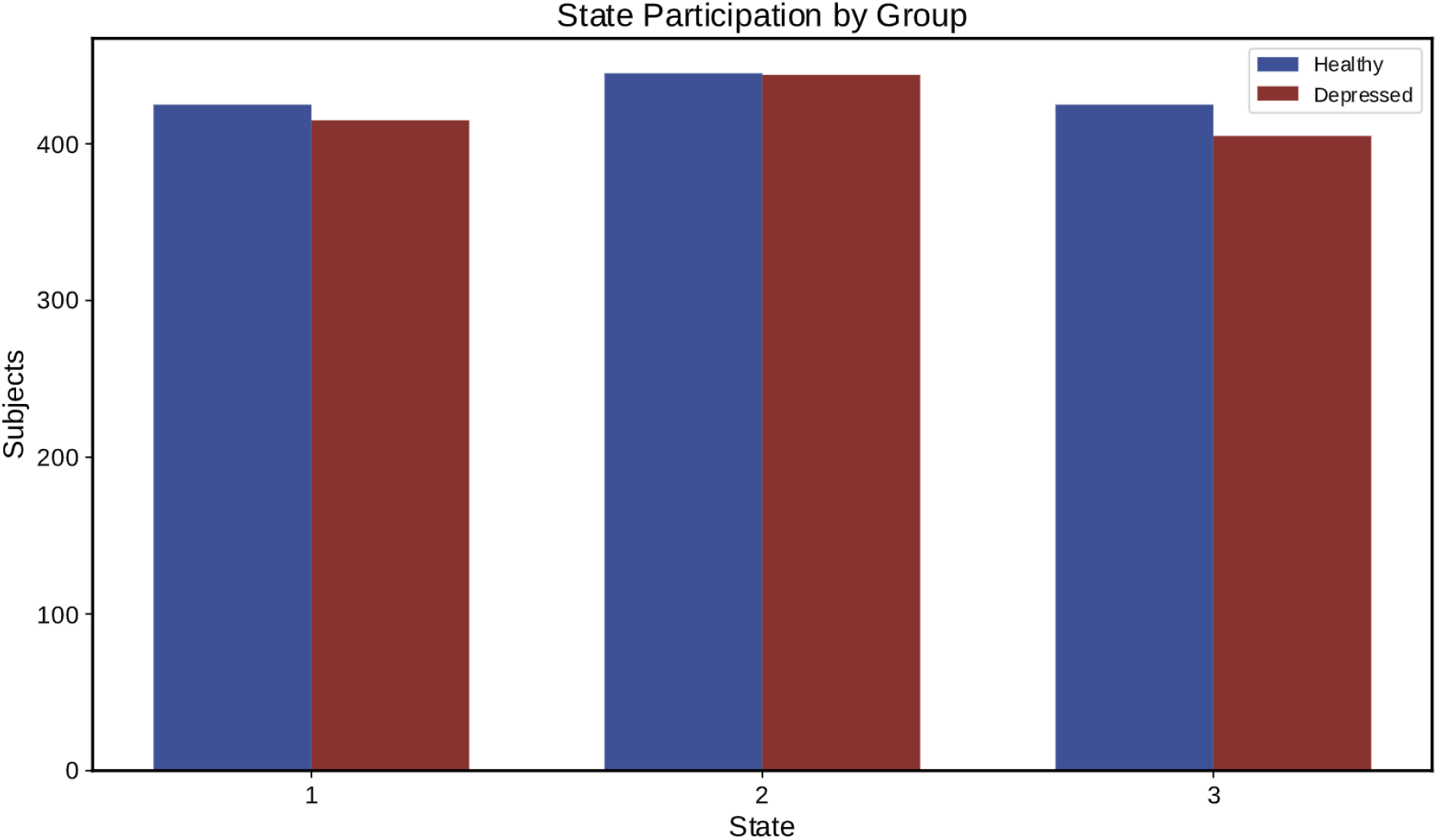
Participant-level engagement with dynamic frontoparietal network (FPN) states across groups. Bars represent the number of participants in each group (patients with major depressive disorder [MDD] vs. healthy controls [HCs]) who occupied each state at least once during the resting-state fMRI scan. The similar distributions support the validity of group comparisons in subsequent temporal dFC analyses.

#### 2.5.2 Between-group statistical comparisons

To test whether dynamic connectivity metrics differed between healthy controls and depressed patients, we conducted independent two-sample Welch’s t-tests for each measure. Group comparisons were performed for fractional occupancy and mean dwell time of each connectivity state, as well as for the total number of state transitions per subject. The Welch’s t-test was chosen, given the heterogeneity of variance between groups. Resulting p-values were adjusted for multiple comparisons using the Benjamini–Hochberg false discovery rate (FDR) procedure (*α* = 0.05).

In a complementary analysis, we further examined differences in the frequency of specific state-to-state transitions between groups. For each subject, we extracted the number of transitions between all pairs of distinct states (self-transitions excluded). These transition counts were compared between groups using Welch’s t-tests, with FDR correction applied across all transitions. This analysis enabled the identification of specific reconfiguration pathways that differentiated patients from controls, beyond differences in overall transition frequency. Effect sizes for all comparisons were quantified using Hedges’ g.

## 3 RESULTS

### 3.1 Demographic and clinical characteristics

The final sample comprised 445 healthy controls (HCs) and 442 patients with MDD. Groups did not differ significantly in age (HC: 36.11 *±* 14.82 years; MDD: 36.47 *±* 12.77 years; *t*(867.7) = 0.38, *p* = 0.698) or sex distribution. Clinical data in the MDD group indicated a mean HAMD-17 score of 21.24 *±* 5.76 and a mean illness duration of 55.65 *±* 73.49 months (Table 1).

### 3.2 Identification of Dynamic Connectivity States

Using k-means clustering applied to windowed FPN connectivity matrices, we identified three recurring dFC states (Figure 4). The optimal number of clusters (*k* = 3) was determined via the elbow criterion (Figure 2). The three states exhibited distinct patterns of functional coupling (Figure 4). State 1 was characterized by moderate connectivity across FPN regions, State 2 by globally reduced connectivity (hypoconnectivity), and State 3 by strongly positive connections.

**FIGURE 4:**
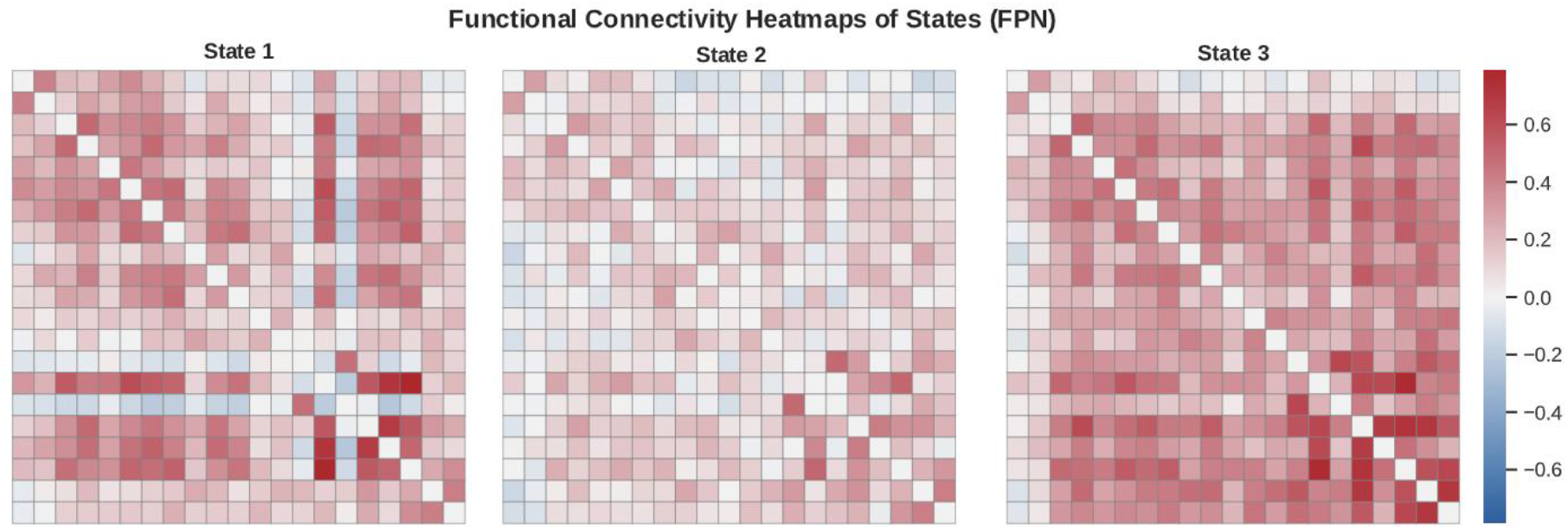
Connectivity matrices for the three dynamic frontoparietal network (FPN) states identified via *k*-means clustering of resting-state fMRI data. State 1 shows moderate functional coupling, State 2 exhibits widespread hypoconnectivity, and State 3 displays strong positive connectivity, especially among lateral prefrontal and parietal regions. Values represent Fisher *z*-transformed Pearson correlation coefficients across 21 FPN nodes from the Dosenbach atlas.

### 3.3 Within-Subject Differences in State-wise FC Strength

To establish whether the identified states represented distinct functional configurations rather than trivial fluctuations, we compared mean FC strength across states within individuals. A repeated-measures ANOVA revealed a highly significant main effect of state, F(1.80, 1597.20) = 5179.65, p < .001 (Greenhouse–Geisser corrected). Greenhouse–Geisser *ϵ* = 0.90). Post-hoc paired comparisons showed that all three states differed significantly from each other after FDR correction (all *q* < 0.001; Figure 5; Table 3).

**TABLE 3:** Within-subject comparison of mean functional connectivity (FC) strength across the three dynamic frontoparietal network (FPN) states. (a) Descriptive statistics (mean *±* standard deviation). (b) Repeated-measures ANOVA results (Greenhouse-Geisser corrected where applicable). (c) Post-hoc paired *t*-test comparisons (Benjamini-Hochberg FDR-corrected). \*\*\**q* < 0.001. FC values are Fisher *z*-transformed Pearson correlations.

| (a) Descriptive Statistics |  |  |  | (b) Repeated-Measures ANOVA |  |  |  |
| --- | --- | --- | --- | --- | --- | --- | --- |
| State | n | Mean | SD | Effect | F | p | $\epsilon$ |
| FC strength State 1 | 836 | 0.198 | 0.043 | State | 5179.650 | 0.000 | 0.902 |
| FC strength State 2 | 886 | 0.106 | 0.025 |  |  |  |  |
| FC strength State 3 | 826 | 0.288 | 0.056 |  |  |  |  |

**TABLE 3** Within-subject comparison of mean functional connectivity (FC) strength across the three dynamic frontoparietal network (FPN) states.
| (c) Post-hoc Comparisons (FDR-Corrected) |  |  |  |  |  |  |
| --- | --- | --- | --- | --- | --- | --- |
| Contrast | n | t | df | p | dz | Sig |
| State 1 vs State 2 | 1605 | 80.25 | 1604 | 0.000 | 2.003 | *** |
| State 1 vs State 3 | 1504 | -57.46 | 1503 | 0.000 | -1.482 | *** |
| State 2 vs State 3 | 1589 | -124.48 | 1588 | 0.000 | -3.123 | *** |

**FIGURE 5:**
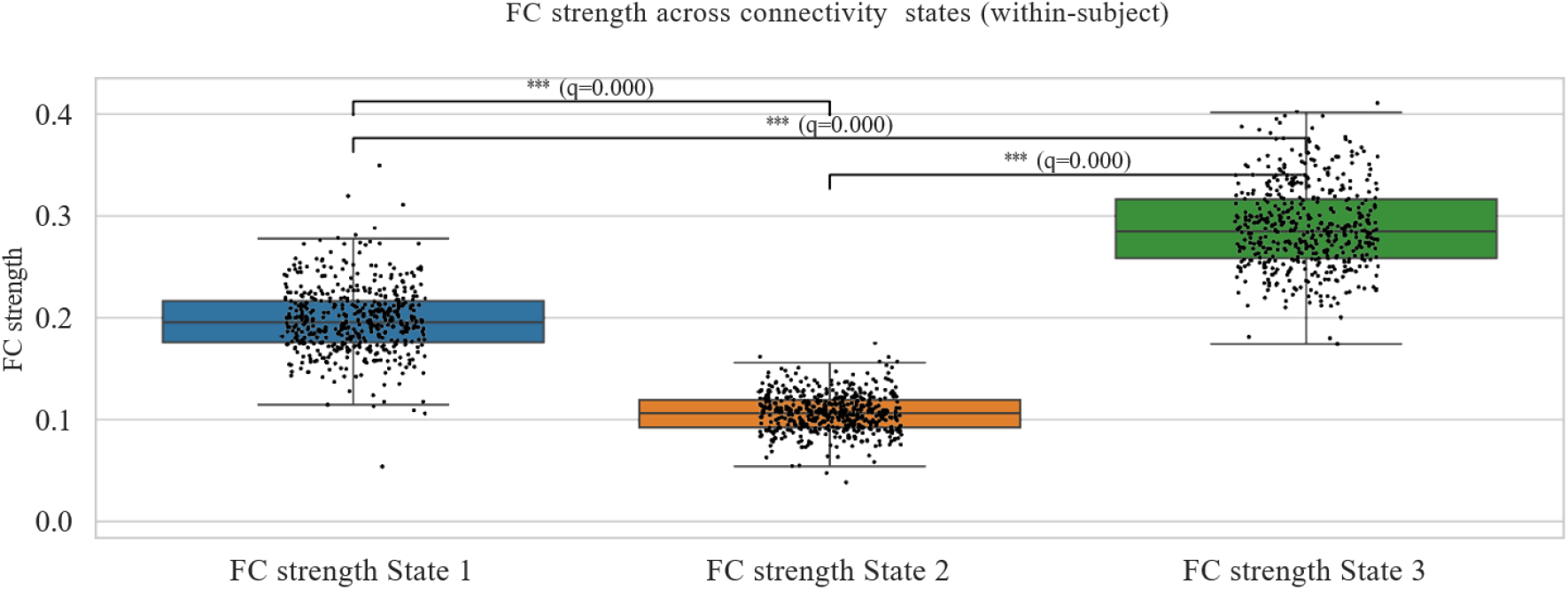
Within-subject comparisons of mean functional connectivity (FC) strength across the three dynamic frontoparietal network (FPN) states. Boxplots show the distribution of Fisher *z*-transformed average pairwise correlations among 21 FPN nodes. State 2 exhibits significantly lower FC (hypoconnectivity), State 3 shows the highest FC, and State 1 is intermediate. Error bars represent the standard error of the mean. Differences were assessed via repeated-measures ANOVA with post-hoc *t*-tests (Benjamini-Hochberg FDR-corrected).

Specifically, State 2 demonstrated the lowest mean FC strength, consistent with a hypoconnected profile, whereas State 3 exhibited the strongest connectivity, and State 1 showed intermediate values. These findings supports that the clustering procedure identified meaningfully distinct dFC states.

### 3.4 Between-Group Comparisons of dFC Temporal Features

We next examined whether the temporal properties of these states differed between groups. Welch’s *t*-tests revealed significant group effects for three metrics (Figure 6 for all metrics; for transitions focus; Table 4). Patients with MDD showed greater fractional occupancy of State 2 (depressed: *M* = 0.553, *SD* = 0.240; healthy: *M* = 0.513, *SD* = 0.219; *t*(876.6) = –2.59, *p* = 0.010, *q* = 0.023, *g* = 0.18), indicating that they spent a greater proportion of time in the hypoconnected state. Patients also exhibited longer mean dwell time in State 2 (*M* = 18.44, *SD* = 27.11) relative to controls (*M* = 13.43, *SD* = 18.15; *t*(769.6) = –3.23, *p* = 0.001, *q* = 0.004, *g* = 0.22), indicating prolonged stabilization within the hypoconnected state. Additionally, the total number of transitions between states was lower in the depressed group (*M* = 22.63, *SD* = 9.33) compared with controls (*M* = 24.84, *SD* = 8.45; *t*(875.3) = 3.69, *p* < 0.001, *q* = 0.002, *g* = 0.25). No significant between-group differences emerged for State 1 or State 3 in either occupancy or dwell time (all *q* > 0.05).

**TABLE 4:**
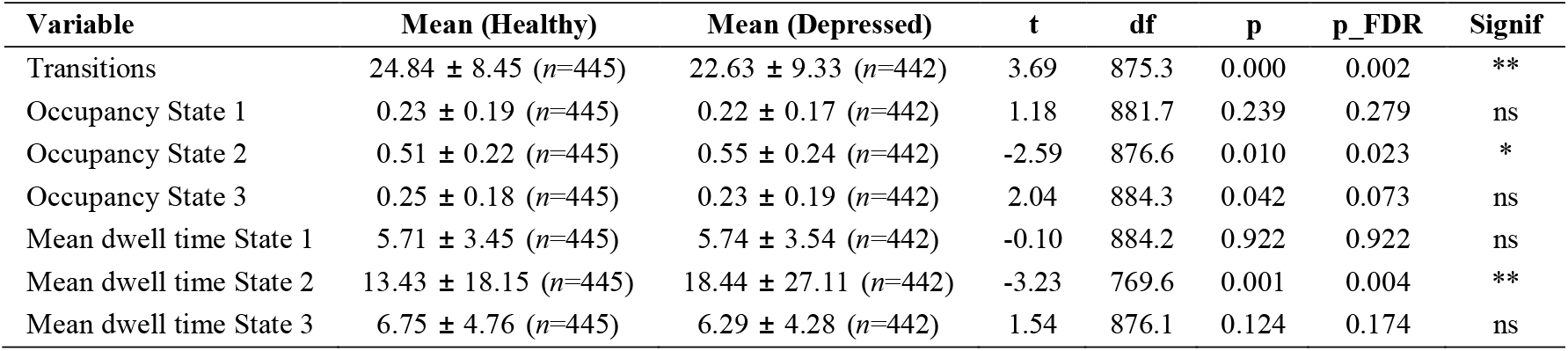
Between-group comparisons of dynamic functional connectivity (dFC) metrics in patients with major depressive disorder (MDD) and healthy controls (HCs). Data are presented as mean *±* standard deviation (with sample size in parentheses). Group differences were assessed using Welch’s *t*-tests, with Benjamini-Hochberg FDR correction applied across comparisons. \*\**q* < 0.01, \**q* < 0.05, ns = not significant.

| Variable | Mean (Healthy) | Mean (Depressed) | $t$ | df | $p$ | $p\_FDR$ | Signif |
| --- | --- | --- | --- | --- | --- | --- | --- |
| Transitions | 24.84 ± 8.45 ($n=445$) | 22.63 ± 9.33 ($n=442$) | 3.69 | 875.3 | 0.000 | 0.002 | ** |
| Occupancy State 1 | 0.23 ± 0.19 ($n=445$) | 0.22 ± 0.17 ($n=442$) | 1.18 | 881.7 | 0.239 | 0.279 | ns |
| Occupancy State 2 | 0.51 ± 0.22 ($n=445$) | 0.55 ± 0.24 ($n=442$) | -2.59 | 876.6 | 0.010 | 0.023 | * |
| Occupancy State 3 | 0.25 ± 0.18 ($n=445$) | 0.23 ± 0.19 ($n=442$) | 2.04 | 884.3 | 0.042 | 0.073 | ns |
| Mean dwell time State 1 | 5.71 ± 3.45 ($n=445$) | 5.74 ± 3.54 ($n=442$) | -0.10 | 884.2 | 0.922 | 0.922 | ns |
| Mean dwell time State 2 | 13.43 ± 18.15 ($n=445$) | 18.44 ± 27.11 ($n=442$) | -3.23 | 769.6 | 0.001 | 0.004 | ** |
| Mean dwell time State 3 | 6.75 ± 4.76 ($n=445$) | 6.29 ± 4.28 ($n=442$) | 1.54 | 876.1 | 0.124 | 0.174 | ns |

**FIGURE 6:**
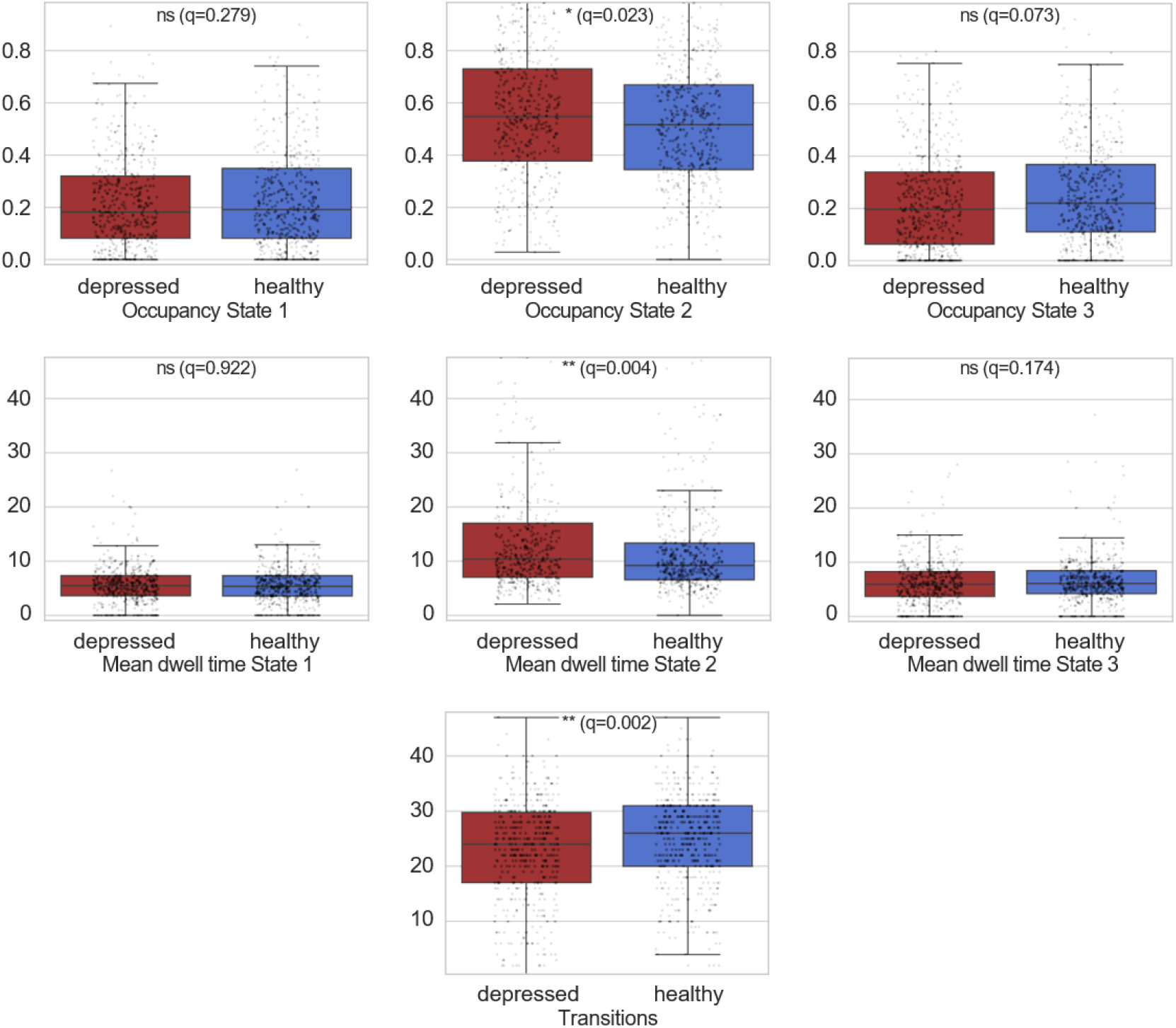
Detailed group comparisons of dynamic functional connectivity (dFC) metrics in the frontoparietal network (FPN). Boxplots display fractional occupancy (top row), mean dwell time (middle row), and total number of state transitions (bottom row) for each of the three states, stratified by group (patients with major depressive disorder [MDD] vs. healthy controls [HCs]). Asterisks denote significant differences (Welch’s *t*-tests with Benjamini-Hochberg FDR correction).

**FIGURE 7:**
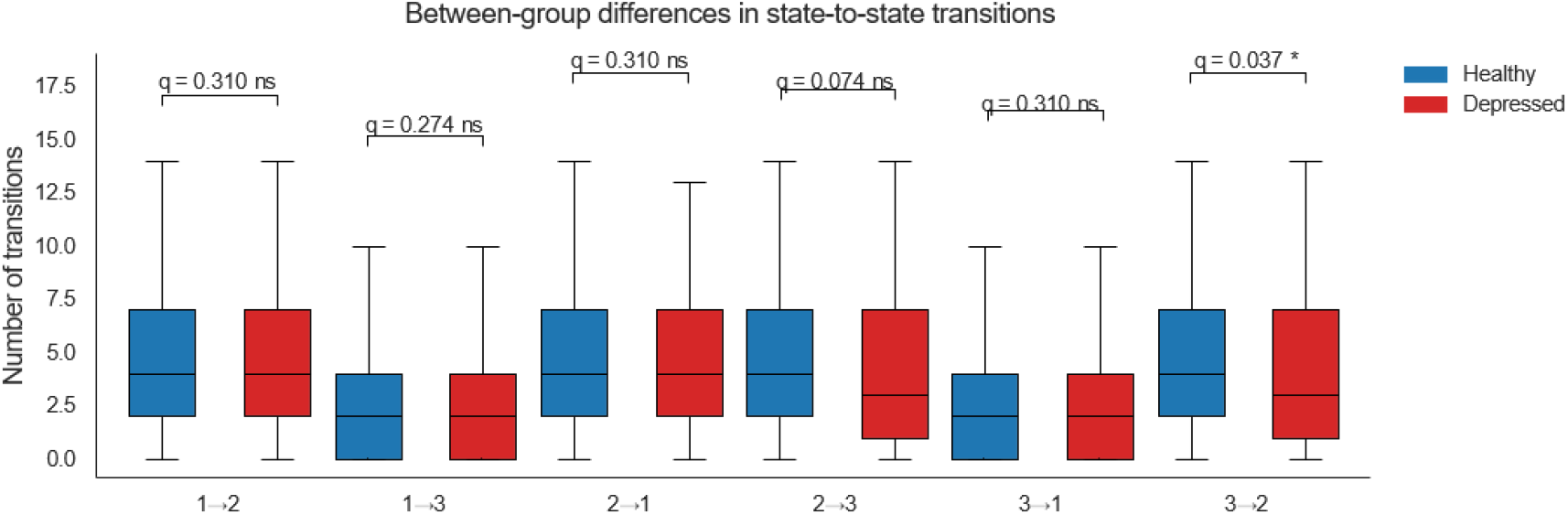
Between-group comparisons of dynamic functional connectivity (dFC) metrics in the frontoparietal network (FPN). Boxplots illustrate differences between patients with major depressive disorder (MDD) and healthy controls (HCs) in fractional occupancy, mean dwell time, and total number of state transitions across the three FPN states. Asterisks indicate significant differences (Welch’s *t*-tests with Benjamini-Hochberg FDR correction).

### 3.5 State-to-State Transition Profiles

To further evaluate the dynamics of switching between states, we compared the frequency of specific state-to-state transitions across groups (Table 5). Welch’s t-tests with FDR correction revealed that patients exhibited significantly fewer transitions from State 3 to State 2 compared with controls (t(885) = 2.75, p = 0.006, q = 0.037). The transition from State 2 to State 3 also showed a nominal group difference before correction (t(877) = 2.25, p = 0.025), although this effect did not remain significant after FDR adjustment (q = 0.074). No other transition pathways approached significance (all q > 0.05). Together, these results suggest a selective disruption in reconfigurations involving State 2, particularly in the ability to shift between the strongly connected state (State 3) and the hypoconnected configuration (State 2), indicating altered flexibility in transitioning between distinct network modes.

**TABLE 5:** Between-group differences in state-to-state transition frequencies in patients with major depressive disorder (MDD) and healthy controls (HCs). Data are presented as mean *±* standard deviation. Group differences were assessed using Welch’s *t*-tests, with Benjamini-Hochberg FDR correction applied across comparisons. \**q* < 0.05; ns = not significant.

| Transition | Mean (HC) | SD (HC) | Mean (MDD) | SD (MDD) | *t* | *p* | *q* | Signif |
| --- | --- | --- | --- | --- | --- | --- | --- | --- |
| 1 → 2 | 4.98 | 3.67 | 4.73 | 3.56 | 1.02 | 0.310 | 0.310 | ns |
| 1 → 3 | 2.64 | 2.70 | 2.37 | 2.73 | 1.49 | 0.137 | 0.274 | ns |
| 2 → 1 | 4.99 | 3.64 | 4.69 | 3.59 | 1.22 | 0.223 | 0.310 | ns |
| 2 → 3 | 4.81 | 3.69 | 4.27 | 3.48 | 2.25 | 0.025 | 0.074 | ns |
| 3 → 1 | 2.64 | 2.71 | 2.45 | 2.79 | 1.03 | 0.304 | 0.310 | ns |
| 3 → 2 | 4.78 | 3.70 | 4.12 | 3.45 | 2.75 | 0.006 | 0.037 | * |

## 4 DISCUSSION

This study presents large-scale evidence indicating that MDD is associated with disruptions in the temporal dynamics of intra FPN connectivity. While previous studies have primarily focused on static functional connectivity (sFC) changes in MDD, our findings extend this line of inquiry and demonstrate that these alterations are also observable in the dynamic organization of brain networks. Using harmonized rs-fMRI data from the REST-meta-MDD consortium (Yan et al., 2019), we identified three recurring FPN connectivity states. It was found that individuals with MDD spend significantly more time in a hypoconnected state, have a longer duration in this configuration, and show an overall reduction in the number of transitions between states. Specifically, a distinct defect was observed in the direct transition between hypoconnected and hyperconnected states. These results indicate that MDD is not only characterized by abnormal connectivity strength but is also associated with reduced temporal flexibility in FPN engagement. This decrease in dynamic adaptability may reflect underlying neural rigidity, which could be a mechanical basis for the cognitive inflexibility commonly observed in depression (Zheng et al., 2024; Menon, 2011). Our first observation was that patients with MDD spent a greater proportion of time in a state characterized by relatively weak coupling among FPN nodes. This finding indicates that hypoconnected configurations of the FPN are more predominant in depression. Because the FPN supports executive functions such as cognitive control, attentional regulation, and top-down modulation (Cole et al., 2014; Harding et al., 2015), greater time spent in a weakly connected state may reduce the network’s capacity to coordinate these processes.

This result is consistent with earlier reports of intra-FPN hypoconnectivity in MDD using static functional connectivity approaches (Kaiser et al., 2015; Hwang et al., 2015; Lan et al., 2022). At the same time, static studies have produced conflicting findings, with some reporting no alterations (Javaheripour et al., 2021; Whitton et al., 2018) and others describing hyperconnectivity in subsets of patients (Zhang et al., 2025; Ren et al., 2025). Such inconsistencies likely arise from the reliance on static functional connectivity, which collapses temporal variability into an average strength measure. In this framework, brief but strong episodes of coupling can disproportionately inflate mean connectivity, masking the more enduring hypoconnected state (Hutchison et al., 2013; Calhoun et al., 2014). By capturing time-varying patterns, the present results highlight the importance of considering dynamic and temporal fluctuations in depression, showing that alterations are not limited to average connectivity strength but extend to how network states evolve and persist over time.

Beyond greater occupancy, patients also remained longer within the hypoconnected state once they entered it. This pattern suggests that transitions out of weakly connected configurations are reduced, leading to prolonged residence in this state. From a dynamic systems perspective, this may reflect the presence of an attractor-like state within the FPN, in which hypoconnected configurations function as stable basins in the network’s dynamic landscape, resisting perturbations and constraining shifts toward more adaptive patterns (Rolls, 2016; Deco et al., 2017).

Prior dFC studies in other large-scale networks have reported similar increases in dwell time in depression, often linked to symptom severity (Demirtaş et al., 2016; Yao et al., 2019). Our results extend this work by demonstrating that the FPN specifically shows this temporal rigidity. Functionally, longer dwell times in hypoconnected states may reduce opportunities for the FPN to shift into more integrated states that could support cognitive control and adaptive regulation. This interpretation aligns with behavioral studies documenting perseveration and impaired set-shifting in depression (Snyder, 2013; Zheng et al., 2024). Nevertheless, it is important to note that our design does not allow us to directly link dwell time to specific symptoms. Future studies correlating FPN dynamics with behavioral or clinical measures will be necessary to clarify these associations.”

A third finding was that patients showed significantly fewer overall transitions between states. Healthy controls exhibited a more diverse temporal profile, frequently alternating between weakly, moderately, and strongly connected configurations, whereas patients showed more restricted switching. This observation parallels prior reports of reduced dynamic range and flexibility of brain states in depression at the whole-brain level (Cole et al., 2013; Han et al., 2020; Wang et al., 2024).

Because transitions are thought to index the brain’s capacity for adaptive network reconfiguration (Bassett et al., 2011; Shine et al., 2016), reduced transitions in depression likely reflect diminished flexibility of the FPN in meeting changing internal or external demands. This reduction suggests a narrowing of the network’s dynamic range, with less frequent shifts among connectivity states of varying strength. Such rigidity may compromise the FPN’s ability to support flexible adjustments in cognitive control, paralleling behavioral evidence of reduced cognitive flexibility in MDD. Interestingly, our findings suggest that major depressive disorder (MDD) is characterized not by a uniform reduction in all frontoparietal network (FPN) dynamics, but by a specific rigidity in transitions between hyper- and hypo-connected states. In healthy individuals, the brain’s metastable organization enables flexible transitions across segregated and integrated network modes, supporting an adaptive balance between stability and flexibility (Deco et al., 2015; Tognoli & Kelso, 2014). This dynamic balance allows the brain to reconfigure connectivity in line with varying cognitive demands, a process tightly linked to effective performance on complex tasks (Shine et al., 2016). By contrast, our results indicate that depressed individuals show a more limited range of brain network configurations, and once their networks enter a hyper-connected state, it is harder for them to directly return to a hypo-connected state (Alonso Martínez et al., 2020; Han et al., 2020).

Such inflexibility at rest may leave individuals less prepared to adjust their cognitive resources when actual demands arise, providing a possible systems-level explanation for the difficulties in attentional shifting and disengagement observed in depression (Koster et al., 2017; Niu et al., 2022). This rigidity may impair rapid switching of attentional focus, consistent with behavioral evidence of deficits in attention shifting and slowed reaction times in depression (Joormann & D’Avanzato, 2010; Snyder, 2013). In short, our findings refine the picture from a global loss of dynamism to a targeted impairment in reconfiguring between opposing FPN regimes.

In sum, this study highlights that major depressive disorder is not only associated with altered connectivity strength but also with disruptions in the temporal dynamics of the frontoparietal control network. Individuals with depression tend to remain longer in hypoconnected states, show reduced flexibility in shifting between configurations, and exhibit selective impairments in direct transitions between distinct states. These temporal alterations provide a mechanistic account of the cognitive rigidity and perseverative tendencies often observed in depression and help to reconcile inconsistencies in prior static connectivity studies. By emphasizing the dynamic nature of frontoparietal dysfunction, our findings underscore the importance of moving beyond static approaches to capture the temporal features of brain network organization in psychiatric disorders.

## 5 CONCLUSION

This study provides large-scale evidence that major depressive disorder is characterized by disruption in the temporal dynamics of the frontoparietal control network (FPN). We identified three recurring connectivity states and found that individuals with depression spend more time in a low-connectivity (hypoconnected) configuration, remain in this state for longer, and generally have fewer transitions between states. The important point is that we observed not all transitions were affected equally; direct transitions between the most different states, i.e., hypoconnected and hyperconnected configurations, were selectively reduced. This pattern shows that the depressed brain not only tends to remain in states with weak connectivity but also has a limited ability to effectively reconfigure across the full spectrum of network dynamics.

These changes indicate a narrowing of the network’s temporal scope and provide a mechanistic explanation for the neural rigidity and cognitive inflexibility observed in depression. By adopting a dynamic perspective, our findings explain the inconsistencies present in static connectivity research and demonstrate that although high-connectivity states exist, they are less accessible, and the dominant feature in depression is a reduced capacity for flexible transitions between states. In summary, this research highlights the importance of temporal dynamics in understanding FPN dysfunction and proposes dynamic connectivity measures as promising options for developing biomarkers in major depressive disorder.

However, the study has some limitations. First, although harmonization between sites was performed, the residual effects of differences in scanners and imaging protocols cannot be completely eliminated. Second, dynamic functional connectivity measures are dependent on methodological choices such as window length, brain parcellation method, and clustering algorithm, which may influence the identified configurations. Finally, we did not examine clinical subgroups of major depressive disorder, such as first-episode versus recurrent cases or treated versus drug-naïve patients; analysis of these subgroups could help determine whether temporal abnormalities represent stable traits, transient states, or treatment-induced changes.

## 6 FUTURE DIRECTIONS

Future research should expand these findings in several important directions. Classified analyses of MDD subtypes will be essential to clarify the heterogeneity in temporal dynamics and to examine whether specific patterns can distinguish different clinical profiles from each other. Linking the characteristics of neurological conditions with behavioral and cognitive measures can also enhance our understanding of how temporal difficulty leads to real-life disabilities. The application of machine learning approaches may also further reveal the value of these temporal indicators for automated classification, prediction, and patient differentiation. Beyond the frontoparietal control network, future studies should analyze not only within-network dynamics but also their relationship to inter-network connectivity, which could illuminate how altered temporal patterns in the FPN interact with broader brain systems. The sum of these steps will refine our understanding of the mechanism of depression and expand the transferability of dynamic connectivity features as biomarkers in clinical research.

## Abbreviations

MDD: Major Depressive Disorder
FPN: Frontoparietal Network
fMRI: Functional Magnetic Resonance Imaging

## ACKNOWLEDGMENTS

We thank the REST-meta-MDD/DIRECT Consortium and the International Big-Data Center for Depression Research, Institute of Psychology, Chinese Academy of Sciences, for providing datasets. We also acknowledge that the dataset providers had no role in study design, data collection, analysis, interpretation, manuscript preparation, or the decision to submit this work for publication.

## CODE AVAILABILITY

The code of this study is available from the corresponding author upon reasonable request.

